# Fetal imidacloprid causes ASD-like impairment of biological motion perception in neonatal chicks

**DOI:** 10.1101/2022.05.19.492744

**Authors:** Toshiya Matsushima, Momoko Miura, Nina Patzke, Noriyuki Toji, Kazuhiro Wada, Yukiko Ogura, Koichi J. Homma, Paola Sgadò, Giorgio Vallortigara

**Author notes:** **Fundings and contribution of the authors:** The present study was supported by grants funded to T.M. by the Japan Society for Promotion of Science (JSPS, Kakenhi; Grants-in-aid for Scientific Research #18K07351, #19KK0211). TM, PS and GV conceived the study. TM and MM designed the behavioral experiments and performed behavioral experiments. MM and NP performed the brain morphometry and isotropic fractionation analysis. NT, KW and KJH designed and performed molecular experiments. YO and TM made statistical analysis. TM wrote the manuscript. All authors agreed on the publication of the paper in its final form.

## Abstract

Several environmental chemicals are suspected as risk factors for autism spectrum disorder (ASD), including valproic acid (VPA) and pesticides acting on nicotinic acetylcholine receptor (nAChR) if exposed during pregnancy. However, their target processes in fetal neuro-development are unspecified. We report that fetal injection of VPA impaired the imprinting of an artifact object in hatchlings, while the predisposed preference to biological motion (BM) remained intact. Blockade of nAChR acted oppositely, namely, spared imprinting and impaired BM in chicks. Beside ketamine and tubocurarine, significant effects of imidacloprid (a neonicotinoid insecticide) appeared at dose ≤1ppm. Despite the distinct processes, both VPA and nAChR blockade similarly impaired imprinting of biological image composed of point-light animation. Furthermore, both impairments were rescued by post-natal bumetanide treatment, suggesting common pathology underlying the social attachment malformation. Ambient neonicotinoid could hinder adaptive socialization through impaired development of visual perception in early neonates.

## Introduction

Despite heterogeneous diagnostic phenotypes, autism spectrum disorder (ASD) is primarily characterized by impaired social interactions1,2. Visual predisposition to animate objects typically arises early in life3–6, which is hampered in neonates/juveniles with ASD or its familial risk7–10. Along with genetic factors, exposure to environmental chemical agents, such as anticonvulsant (valproic acid, VPA)11–14 and pesticides15–18 during pregnancy is an ASD risk factor. Despite intensive efforts to develop mammalian models to assess these environmental factors19,20, the validity is limited because these neonatal mammals (mostly rodents) do not spontaneously perceive biological motion (BM) in early life, which is found only in human neonates3–5 and a taxonomically distant animal, domestic chicks21.

The ability to visually perceive conspecifics, as well as identify individuals, their emotions and intention by highly reduced stimuli (such as point light animations) is referred to as the BM perception22. Generally, motion perception develops slowly in human neonates with early visual experiences exerting significant influences23. Pre-attentive processing of biologically relevant stimuli (BM and face configuration) canalizes the subsequent memorization, leading to the adaptive formation of social attachment to specific individuals4. Human neonates typically show an early emergence of BM preference3,4, which most non-human animal species do not reveal. Various non-human animals discriminate BM point-light animations24–27, but their BM perception does not spontaneously arise after birth.

In chicks, the spontaneous BM preference21 is enhanced by imprinting of an artifact (non-BM) object28, facilitating and canalizing the subsequent imprinting29,30. Along with the commonality in the early development of BM, humans31 and chicks32 show inversion effects, namely predisposition to up-right walking configuration. Assuming the different brain organizations between birds and humans, the functional and developmental similarities are striking. Although the neural substrates of the neonatal BM perception are largely unspecified, recent studies suggest the sub-cortical visual processing as evolutionarily conserved visual systems33,34.

A question arises regarding this context. What prenatal processes construct the BM preference? Previous studies revealed the effects of fetal VPA application on social behaviors in chicks35–38, but the BM preference has not been addressed. Furthermore, if the VPA effect on the social behavioral hypoplasia was mediated by well-documented action as an inhibitor of histone deacetylation39, what other risk agents did? As VPA is a potent anti-convulsant drug, it might effectively suppress the fetal movements, hence, the BM preference. As the first step, we hypothesized that suppression of spontaneous fetal movements40,41 is responsible, where excitatory GABA actions could play a role42,43. In human adults, execution of an unfamiliar motor patterns facilitates perception of the corresponding BM animations, thus, non-visual motor execution might be involved44. According to the motor involvement, an imaging study suggested contribution of the posterior part of cerebellum in BM perception45. If this applied also to fetuses, any chemical agents that suppress spontaneous movement, including the blockade of neuro-muscular transmission, would impair the development of BM preference. We started this study by listing chemical agents that effectively suppress the fetal movement at embryonic day 14 (E14), the stage wherein VPA is reported to be effective35.

## Results

### Suppression of the spontaneous fetal movement by chemical agents

The fetal movement was reliably detected by placing an analog record stylus against the shell surface (experiment 1, **Figure 1A**; see supplementary for the methods and statistical analyses). Low frequency power (2~20 Hz) of the recorded signal (ballistogram46) revealed that VPA injection to the air sac acutely suppressed the movement (**Figure 1B**). Similar suppression was found by ketamine (sedative drug of a wide action spectrum including blockade of NMDAR and nAChR), mk801 (selective NMDAR blocker), and tubocurarine (nAChR blocker of a wide spectrum) (**Figure 1C**; see supplement for the chemical agents used and statistics). Among other nAChR blockers, methyllycaconitine citrate (MLA, specific to neuronal α7 subtype), and dihydro-β-erythroidine hydrobromide (DHβE, specific to muscular α4β2 subtype) were similarly effective. Fetal movement is sensitive to the blockade of cholinergic or NMDAR-mediated neurotransmission in the brain (VPA, ketamine, mk801, and MLA), and neuro-muscular junctions in the periphery (DHβE); tubocurarine can act centrally and peripherally.

**Figure 1.**
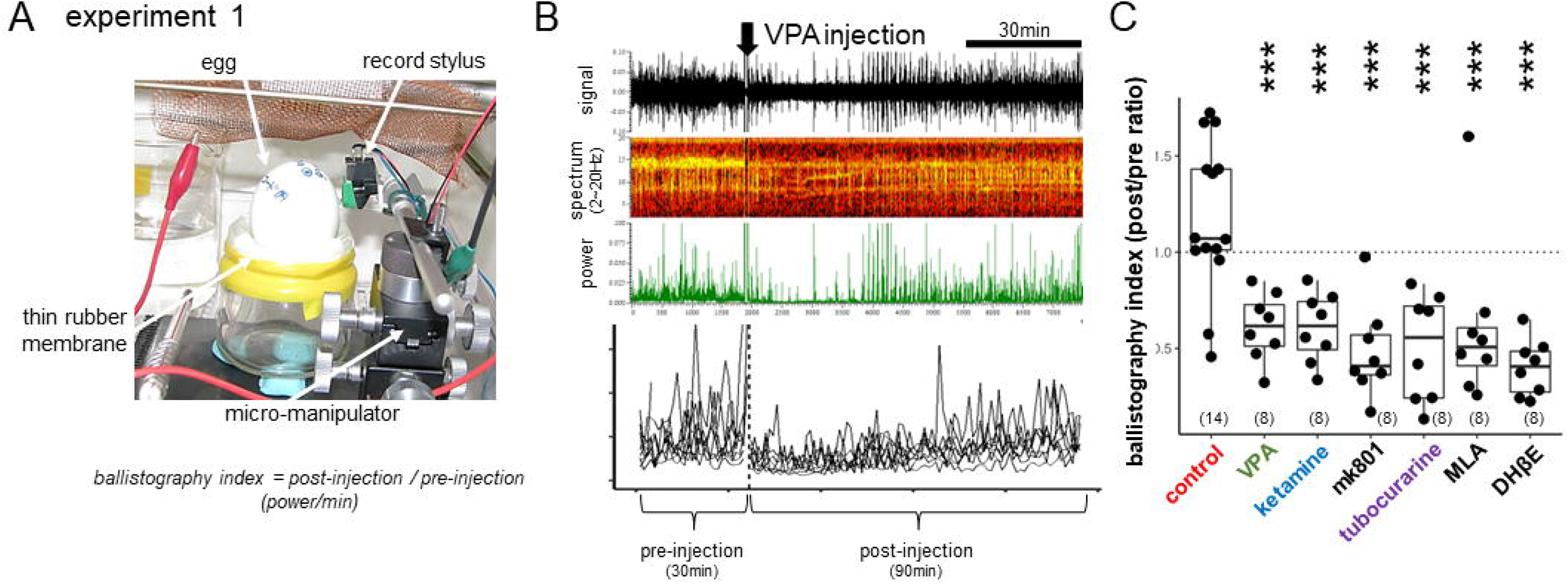
Suppression of the spontaneous fetal movement by chemical agents. **(A, B)** Ballistography recording (experiment 1) and blockade by sodium valproate (VPA). Record stylus placed against the eggshell surface detected the miniature vibration caused by spontaneous fetal movement. **(C)** The dose was experimentally searched for each agent so that fetal movement was similarly suppressed on embryonic day 14 (E14), wherein the treatment gave rise to normal hatch rate and successful training of the hatchlings. Asterisks indicate the significant difference from the control by multiple regression analysis; *, p<0.05, **, p<0.01, ***, p<0.001. Number in parenthesis indicates the sample size.

### BM preference and imprinting were doubly dissociated in chicks

Hatchlings of the injected eggs were tested for BM preference and imprinting (experiment 2). Chicks were individually trained by a video clip of a rotating red toy (displayed on an LCD with sound) for two hours with one hour intermission, and after 30 to 60 min, tested for BM preference (walking point-light animation over linear motion, both in white) and for imprinting memory (familiar red toy over novel yellow) (**Figure 2A**). Among the examined agents, ketamine, tubocurarine and MLA significantly reduced BM preference compared with control in a dose-dependent manner, while VPA failed (**Figure 2Ba**; see **Figure S2 and 3** for details). Despite effective suppression of the fetal movement, mk801 failed to cause any behavioral effects, and hence, VPA and ketamine should act through pathways other than NMDAR. Moreover, nAChR is responsible for the BM preference impairment because ketamine, tubocurarine and MLA were similarly effective. Notably, besides NMDAR suppression, ketamine directly blocks the nAChR-associated channels in the central nervous system and peripheral tissues47–48. However, VPA spared BM and impaired imprinting (**Figure 2Bb**), indicating that the underlying pharmacological mechanisms are doubly dissociated between these two behavioral traits, except that MLA and DHβE weakly suppressed imprinting. Suppression of fetal movements is therefore not sufficient for the BM preference impairment in hatchlings.

**Figure 2.**
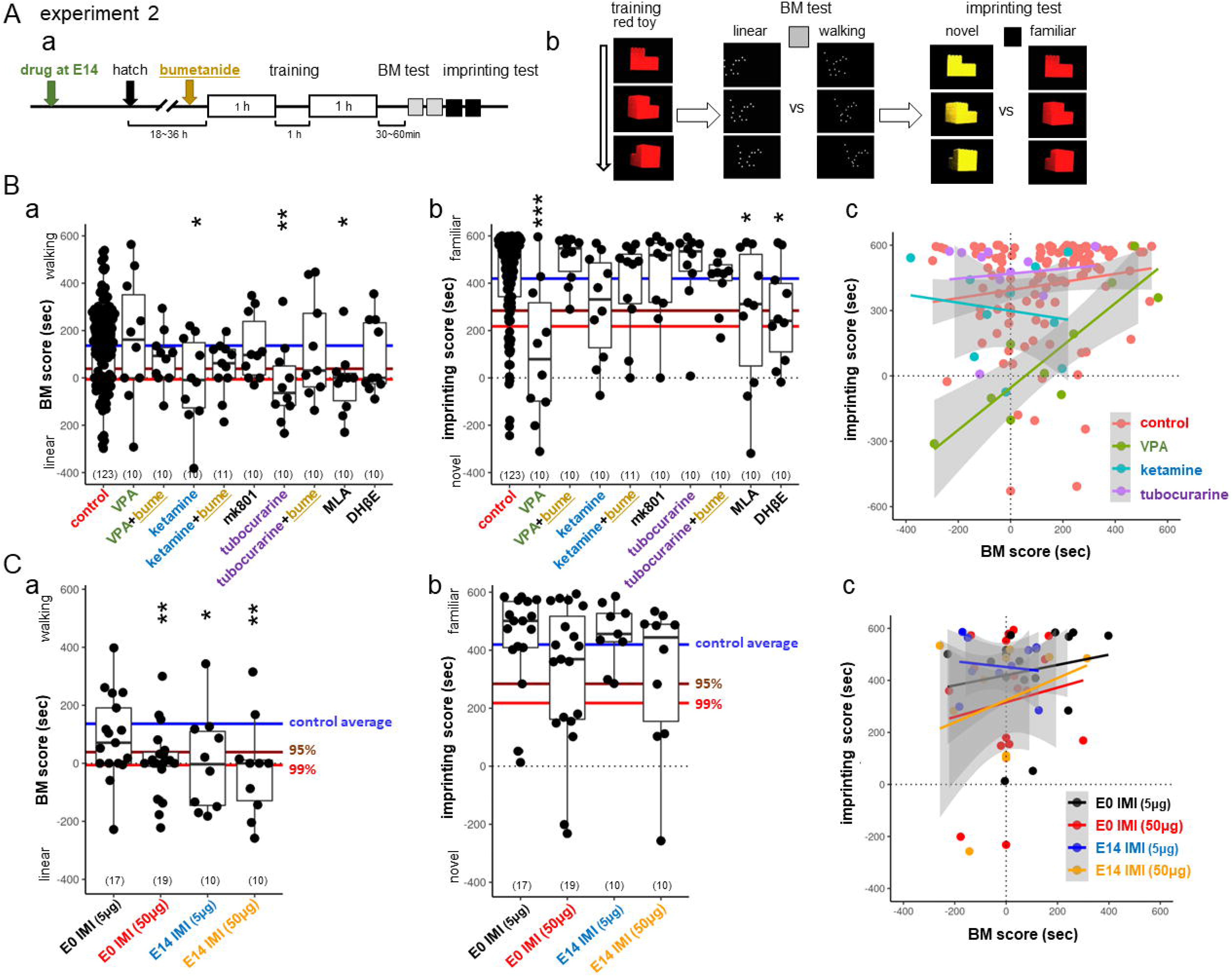
BM preference and imprinting were doubly dissociated in chicks. Effects of the fetal injection of chemical agents on the biological motion (BM) preference and imprinting memory were examined in the hatchlings. **(A)** Training and test procedures (experiment 2). Chicks were trained by an artifact object (rotating red toy) and tested by binary choices for BM (walking motion vs linear motion) and imprinting (red toy vs yellow); stay time difference (sec) was used for the scores. **(B)** BM **(a)** and imprinting score **(b)** of the treated hatchlings were plotted (**c**). Horizontal lines indicate the critical levels of the control data determined by bootstrap computation; average (blue), 95% (brown) and 99% (red) confidence levels. Distinct spectrums were found between VPA and other agents. **(C)** Imidacloprid (IMI) also suppressed BM **(a)** if injected on E0 or E14. Although not significant, imprinting was slightly suppressed by IMI 50μg in both injection days **(b)**.

**Figure 3.**
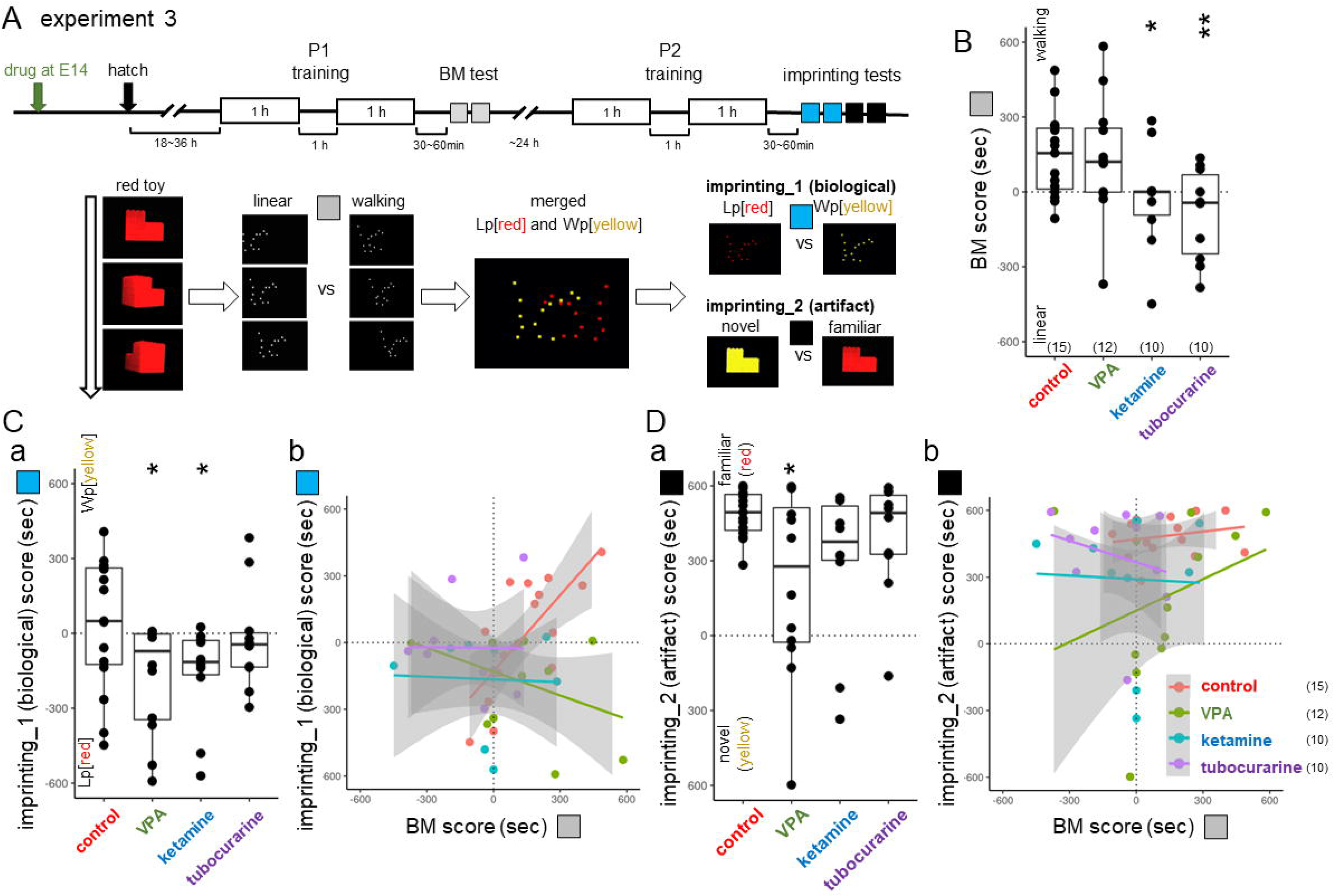
Imprinting of biological motion image was impaired by both VPA and nAChR blockade. The two-step imprinting paradigm revealed common hypoplasia by E14 injection of VPA and nAChR blockade (experiment 3). **(A)** Behavioral procedure. After the post-hatch 1-day (P1) training using an artifact (rotating red toy) and BM test, chicks received the second day (P2) training, wherein two images of point light animations were presented with yellow in walking (Wp[yellow]) and red in linear rigid motion (Lp[red]). **(B)** BM scores confirmed the experiment 2. **(Ca,b)** Both VPA and ketamine chicks showed the impaired formation of preference to Wp[yellow]. Notably, the tubocurarine chicks showed similar plots to the ketamine. (**Da,b**) VPA chicks showed the impaired imprinting to artifact red toy as in experiment 2.

Distinct behavioral phenotypes appeared when imprinting score was plotted against BM score (**Figure 2Bc**). The control chicks showed a weak but significant positive correlation between the two scores (r= +0.21, p=0.0217 for n=123, Spearman’s rank correlation), which was stronger in the VPA chicks (r= +0.88, p= 0.000747 for n=10) but absent in the ketamine (r= −0.042, p=0.919) and the tubocurarine chicks (r= −0.10, p=0.785). Although the cause of the individual variations was unspecified, those chicks with high BM score were resistant to VPA applied on E14. On the other hand, ketamine and tubocurarine spared the imprinting score in these chicks with low BM scores.

Blockade of nAChR transmission by imidacloprid (IMI) also impaired BM in a dose-dependent manner (**Figure 2C**). When applied to E14, IMI suppressed fetal movement at the low dose (5-50 μg/egg: ~0.1-1.0 ppm), and significantly impaired BM but spared imprinting in hatchlings. As mammalian fetus could be maternally exposed to ambient neonicotinoids, we tested the IMI effect injected into fertilized eggs before incubation (E0). The treated hatchlings showed a significant impairment of BM at 50 μg/egg. Plotting the imprinting score against the BM (**Figure 2Cc**) revealed that the chicks with a low BM score tended to show a lower imprint score in the high-dose IMI groups (50 μg/egg), though without significant correlations (r= +0.24 and p=0.314 for E0, r= +0.18 and p= 0.623 for E14).

### Bumetanide rescued both of the impairments by VPA and nAChR blockade; shared pathology at the molecular level

In concert with the excitatory nature of GABA transmission in fetus43, injection of bumetanide (blocker of a chloride cotransporter, NKCC1) on E14 also impaired the post-hatch BM preference, whereas imprinting was spared (see Supplementary material, **Figure S2C,D**), suggesting a critical importance of excitation/inhibition balance during the late phase of fetal development. When applied to P1 chicks (30 min before the start of the training session; 0.02mg/chick), on the other hand, bumetanide rescued both impairments, namely imprinting (for VPA) and BM (for ketamine and tubocurarine; **Figure 2Ba,b**). Despite distinct pathogenesis, BM and imprinting impairments could share common molecular bases associated with GABA-AR actions. Bumetanide and VU0463271 (KCC2 blocker) had no significant effects on the spontaneous movements of E14 fetuses (**Figure S1B**); suppression of the fetal movements is not necessary for the BM impairment.

### Imprinting of biological motion image was impaired by both VPA and nAChR blockade; shared behavioral phenotype

BM preference canalizes the subsequent imprinting31. If chicks were trained by a non-biological artifact (rotating toy in red) on the post-hatch one day (P1), they were oriented to a biological object (walking point in yellow, Wp[yellow]) on the subsequent day (P2; experiment 3, **Figure 3**). Injection of ketamine and tubocurarine (but not VPA) on E14 suppressed BM preference on P1 (**Figure 3B**) as in experiment 2. However, the P2 training using merged point-light animations (linear motion in red and walking in yellow, Lp[red]+Wp[yellow]) resulted in the impaired formation of preference for Wp[yellow] in all groups (imprinting_1 score in **Figure 3Ca**), while preference to the artifact (red toy) remained intact except VPA (imprinting_2 score in **Da**). Multiple regression analysis of the imprinting_1 score revealed a significant interaction of BM score and each of VPA, ketamine, and tubocurarine. VPA and nAChR blockade similarly impairs the social attachment formation to biological image via distinct neurodevelopmental processes.

### Distinct effects of VPA on histone acetylation, brain size and neuronal maturation

VPA is shown to act as a potent inhibitor of histone deacetylases (HDACs)39. To see if ketamine (as nAChR blockader) could also modify HDACs, we examined histone acetylation in primary cell culture prepared from E14 embryonic telencephalon; fluorescence level was measured in randomly selected 100 cells for each treatment (experiment 4, **Figure 4A**). As expected, VPA increased the H3K27 acetylation level at a dosage comparable to the *in ovo* injection (0.12~1.2 mM of medium), whereas ketamine had no effects at 6.6 and 66 μg/mL, which was higher than the *in ovo* dose (0.2mg/50g = ca. 4 μg/mL). The VPA effect on HDACs was confirmed, whereas no comparable effects were found by ketamine.

**Figure 4.**
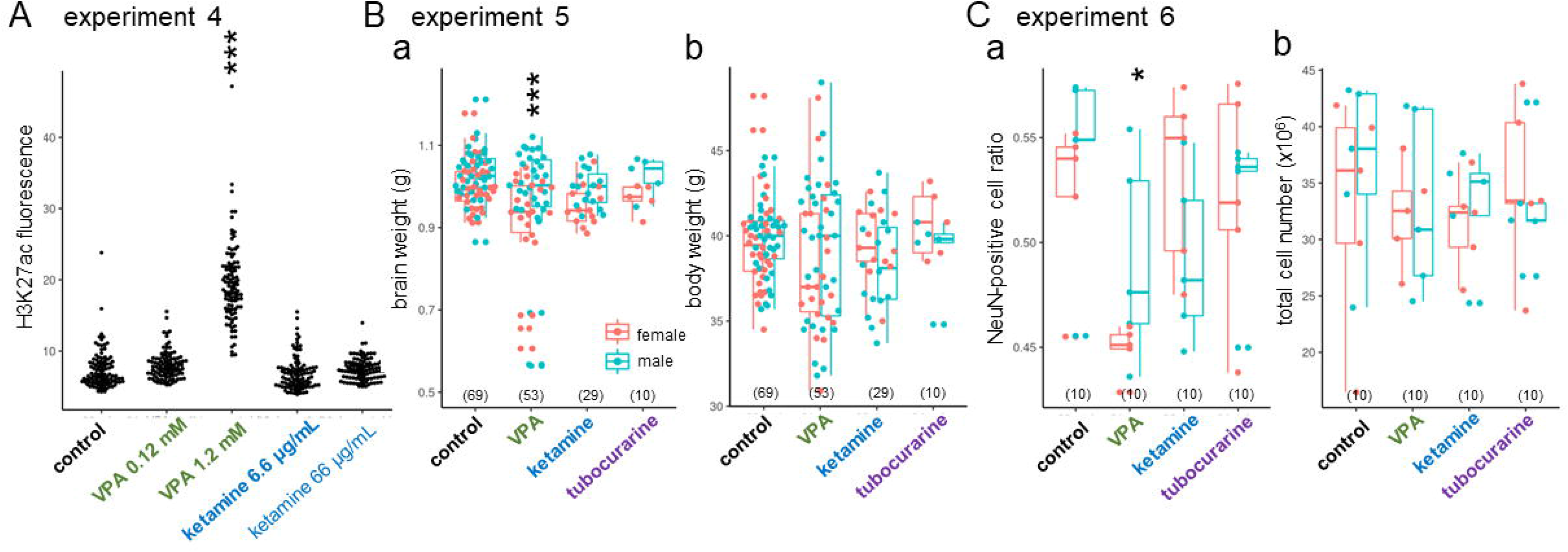
Distinct effects of VPA on brain size, matured neuron ratio, histone acetylation, and expression of chloride co-transporter genes. **(A,B)** VPA but not ketamine/tubocurarine reduced the brain weight **(Aa)**, and the ratio of NeuN positive cells measured by isotropic fractionation **(Bb)**; body weight **(Ab)** and total cell number **(Ba)** were not affected. Male brains were significantly heavier, but no interaction was found between the sex and the agents. No significant effects of sex were detected on the cell number and the NeuN ratio. **(C)** VPA but not ketamine enhanced the acetylation level of histone H3K27 in primary cell culture made from E14 telencephalon. **(D)** Expression of chloride cotransporters (ratio of NKCC1/KCC2 for *slc12a2/slc12a5*; **Cb**) was compared in hatchlings after VPA **(Da)** and tubocurarine **(Db)** injection on E14. Marginally significant interaction (p=0.067) was found in the tectum of VPA chicks.

Different brain morphology could appear in these chicks with fetal chemical treatments. Although controversial, macrocephaly (increased head/brain size) is associated with some ASD subtypes due to altered neurogenesis in the early infantile period49,50. In chicks, the whole-brain weight of hatchlings revealed that E14 injection of VPA, but not ketamine, significantly decreased the brain weight compared to control (experiment 5, **Figure 4Ba**) without changes in the body weight (**Bb**). Notably, 11 out of 53 chicks (~21%) had a distinctly small brain, whereas the rest remained in the control range. Male brains were significantly bigger, but no significant interaction occurred between sex and treatment. Isotropic fractionation revealed a significantly lower ratio of NeuN-positive cells occurred in VPA (**Figure 4Ca**) without changes in the total cell number (**Cb**). Perturbation of gene expression by VPA could lead to retardation of brain development during the late fetal stage and, hence, the delayed maturation of neurons.

In conclusion, suppression of fetal movement is not sufficient or necessary for the social attachment impairment. The present study revealed two critical neurodevelopmental processes, namely the BM predisposition and the memory formation; the former depends on the nAChR transmission, whereas the latter is particularly fragile to VPA. Despite distinct, both processes are critical for hatchlings to form social attachments to biological objects, and share a common pathology of delayed GABA switch when impaired. Finally, imidacloprid impaired both BM and imprinting at the sublethal low dose of ≤1ppm.

## Discussions

### Plausible scenario of the adaptive socialization through imprinting

Since Lorenz51, imprinting has been assumed as a simple but unusual type of learning in limited precocial animals, where irreversible one-time learning occurs in a restricted short critical period after birth. This study series suggest a different figure: imprinting is composed of multiple processes and mechanisms functioning together (**Figure 5**). In the early neonatal processes, exposure to any moving object leads to the CONCPEC mechanism52 associated with the thyroid hormone (specifically, triiodo-thyronine T_3_) influx, which is induced by the *primary imprinting* and, in turn, elongates the sensitive period53 and enhances innate predispositions54,55. Therefore, subsequent *secondary imprinting* is canalized to objects bearing biological features, namely, the CONLERN mechanism is activated toward adaptive social attachment. These early processes, which are homologous to that proposed for neonatal development in humans34.

**Figure 5.**
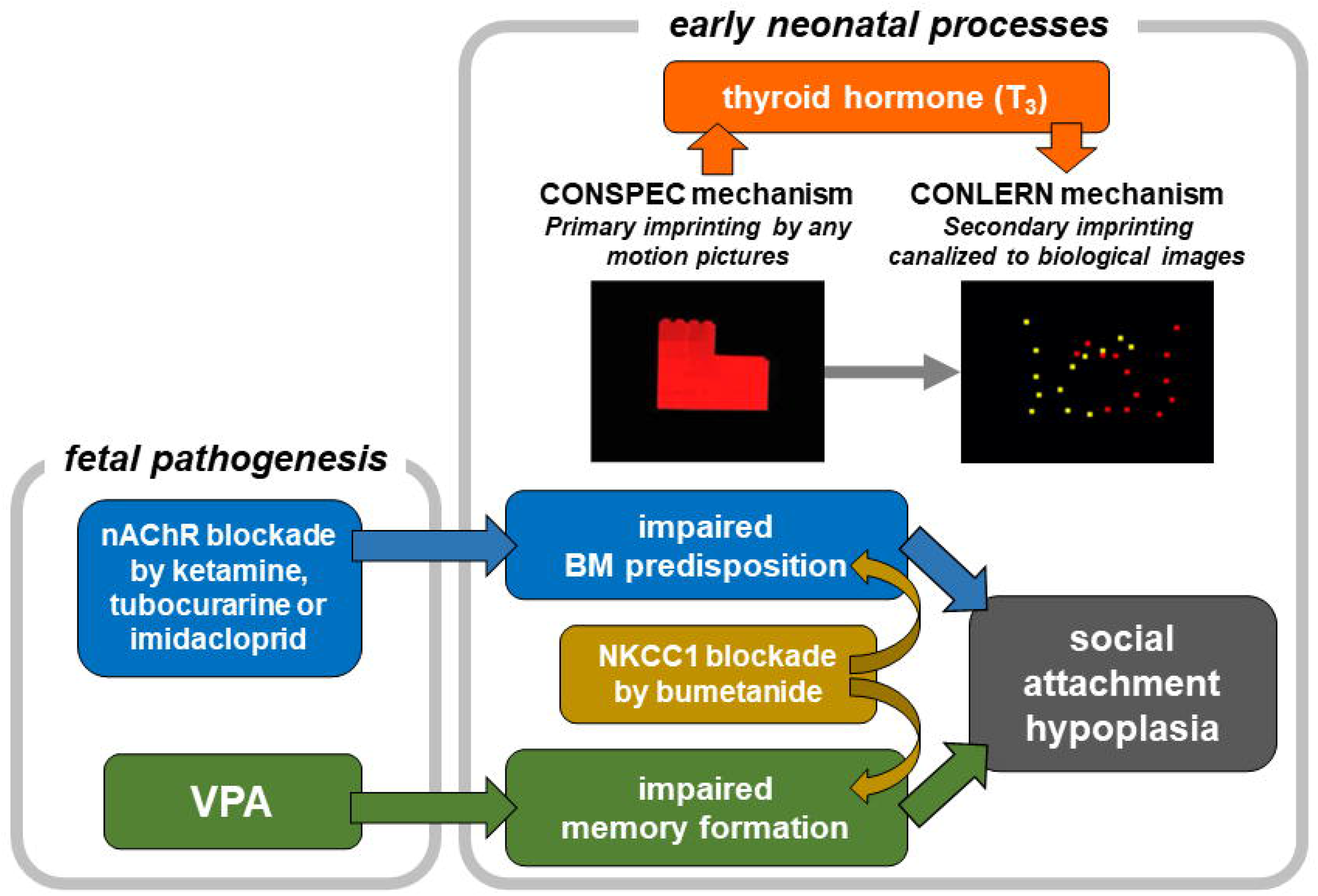
Plausible scenario of the adaptive socialization through imprinting.

### Central and peripheral targets of the nAChR blockade

Nicotinic transmission is critical for neurodevelopment56,57 including the pathophysiology of ASD58,59. However, as nAChR is widespread in the brain and body60,61, several brain regions/organs must be considered. Among the possible candidates, the following regions need specific attention: (1) cholinergic neurons in the striatum and the basal forebrain projecting to wide areas in pallium (or cortex in mammals)62, (2) cholinergic neurons in nucleus isthmi parvocellularis (or pedunculo-pontine tegmental nucleus) projecting to optic tectum (superior colliculus)63, and (3) pre-ganglionic cholinergic terminals of parasympathetic nerve acting on thyroid gland64. If (1) is the case, involvement of telencephalon in the BM predisposition should be considered as delayed GABA switch may be an underlying factor in the brain areas along with IMM, arco and tectum. If (2), retarded visual attention due to the underdeveloped tecto-isthmo-tectal network must be considered. If (3), hypothyroidism during gestation or early neonatal period could be the target as reported in human neonates65,66 and chicks53. The responsible regions’ specification will enable us to design appropriate amendments for each phenotype.

### Neonicotinoids could cause ASD via neonatal vision

Environmental risk of neonicotinoids was initially related to the population decline of insectivorous birds67 and the delayed migration by impaired foraging behavior68. Recently, the association between ambient neonicotinoids and ASD in humans has been identified in several extensive epidemiological studies69–71. Fetal/neonatal exposure of acetamiprid neonicotinoid caused abnormal development of socio-sexual behavior in mice72. Despite the low level of acute toxicity, low affinity for vertebrate nAChR, and rapid metabolism73, neonicotinoids are known for a high persistence level. As shown in this study, identical effective dosage appeared in the E0 and E14 groups of IMI treatment (**Figure 2C**). IMI suppressed BM preference nearly completely at 1ppm (1mg/kg egg weight), which was significantly lower than the lethal dose (LD_50_ <44 mg/kg/day in red-legged partridges) and even lower than the dose (8.8mg/kg/day) that reduced immune response of the offspring74. Legislation on the control of residual neonicotinoids and other pesticidal chemicals in feed, meat, and eggs should consider the risk of increase in the ASD cases by nicotinic blockers.

### Advantages and limitations of chicks for screening of ASD risk agents

This study suggests that the domestic chick is a valid animal model75,76 for studying ASD, even though birds are taxonomically distant from humans. (1) Chick has construct validity because there are common causes and processes, such as VPA, blockade of nAChR, and involvement of GABA transmission. (2) Chick uniquely has face validity because BM preference and learned social attachment formation are assayed. (3) Chick has predictive validity because bumetanide rescued the impairments by VPA and nAChR blockade. We must also consider the recent development of a rich repertoire of behavioral paradigms available in domestic chicks77–79, ranging from numerical comprehension, arithmetic, and the link of the social rank to cognitive capability. ASD is characterized by the impaired social interactions and other symptoms such as delayed speech, learning disability, repetitive stereotyped behaviors, and altered (reduced or sensitized) sensory perception, which we may address using chicks as the model, including a longitudinal life-long survey of these treated chicks.

Although inferior in the homological validity76, the chick model has several technical advantages over conventional mammalian models. (1) Maternal complication is disregarded, whereas gestational effects are inevitably subjected to the maternal metabolism in mammals. (2) The time course and effective dose of agents are precisely determined. (3) The short incubation period facilitates speedy screening, as it takes only a week from the injection of chemicals to E14 eggs until the hatchling are tested on P1. (4) Simple imprinting paradigm allows testing of predisposed preference and learning ability. Humans are often exposed to several ASD risk factors simultaneously. The chick model is suitable for examining interactions of these agents in a strictly controlled manner by monitoring heterogeneous aspects of the disabilities.

## Acknowledgements

Critical comments and instructive suggestions by Dr. Brian McCabe (University of Cambridge) are highly acknowledged. Contribution of Mr. Yasutaka Sasaki (Machine Department of the Faculty of Science, Hokkaido University) must be acknowledged for production of the imprinting apparatus. We also thank Editage (www.editage.com) for English language editing.

## Supplementary materials: Methods and Statistical Analyses

### 1. Methods

#### 1.1. Animals and compliance with ethical standard

Experiments were conducted under the guidelines and approval of the Committee on Animal Experiments of Hokkaido University (approval number 20-0141). The guidelines are based on the national regulations for animal welfare in Japan (Law for Humane Treatment and Management of Animals, after partial amendment No. 68, 2005). Number of subjects used are indicated below in the statistical analyses for each experiment.

Domestic white Leghorn chicks (*Gallus gallus domesticus*, an egg-laying strain of Leghorn called “Julia”) were used. Fertilized eggs were purchased from a local hatchery (Iwamura Co., Niigata/Hokkaido Japan) every 2 weeks, and the batch was numbered. Eggs were incubated in the laboratory by using type P-008B incubators (Showa Furanki Co., Saitama Japan) with its temperature controlled at 37.7°C and the humidity at ca. 80%. The inside of the incubator was kept in complete darkness until hatch. Chicks were individually hatched in small boxes separated by black plastic walls, so that they could interact acoustically but not visually. To avoid post-hatch visual experiences, hatchlings were individually housed in boxes placed in another incubator of the same type, kept in darkness until the experiment. Chicks were sexed after the experiment. Generally, fertilized eggs weighed 59.8 ± 3.1 g (mean ± s.d., n=170) and ca. 2 g lighter at embryonic 14 days (E14) whereas the shells weighed 8.0 ± 0.1 g (n=16), so that the wet weight is assumed to be around 50 g. In this series of experiments, as the egg weight did not considerably vary, the amount of injected chemical per egg was fixed and not adjusted for each egg.

#### 1.2. Ballistographic recording of embryonic movements (experiment 1)

Mechanical vibration caused by spontaneous fetal movements were recorded following the ballistocardiogram method developed by Suzuki et al. (1989; reference #47). By a micromanipulator, an analog record stylus cartridge (type AT VM95E, Audio Technica Co., Tokyo Japan) was gently pushed against the shell of egg placed on a thin rubber membrane, which was stretched on a glass jar (**Figure 1A**). The jar and the manipulator were firmly settled on a heavy iron plate and housed in a constant temperature incubator at 37.7°C and high humidity. The recorded monoaural signal was amplified by a hand-made low input-impedance amplifier (made of FET operational amplifier, TL084), band-pass filtered (cut-off frequency: 10~100Hz, magnification: x1,000), and stored at 1,000 point/sec sampling rate by Spike2 (ver.7, the interface CED-1401 micro3, Cambridge Electric Design Co., Cambridge UK). Frequency spectrum of 2-20 Hz range was monitored, and the converted power was stored at every 1 sec (**Figure 1B**). Using R as platform (see below), the power value was averaged at every 1 min, and the temporal profiles were constructed for pre-injection 30 min and post-injection 90 min for each egg.

Superimposed traces of the ballistographic recordings obtained from 9 sets of fetuses are shown in **Figure S1A**. Dose dependency of the suppressive effects of injected chemicals are shown in **Figure S1B**.

**Figure S1.**
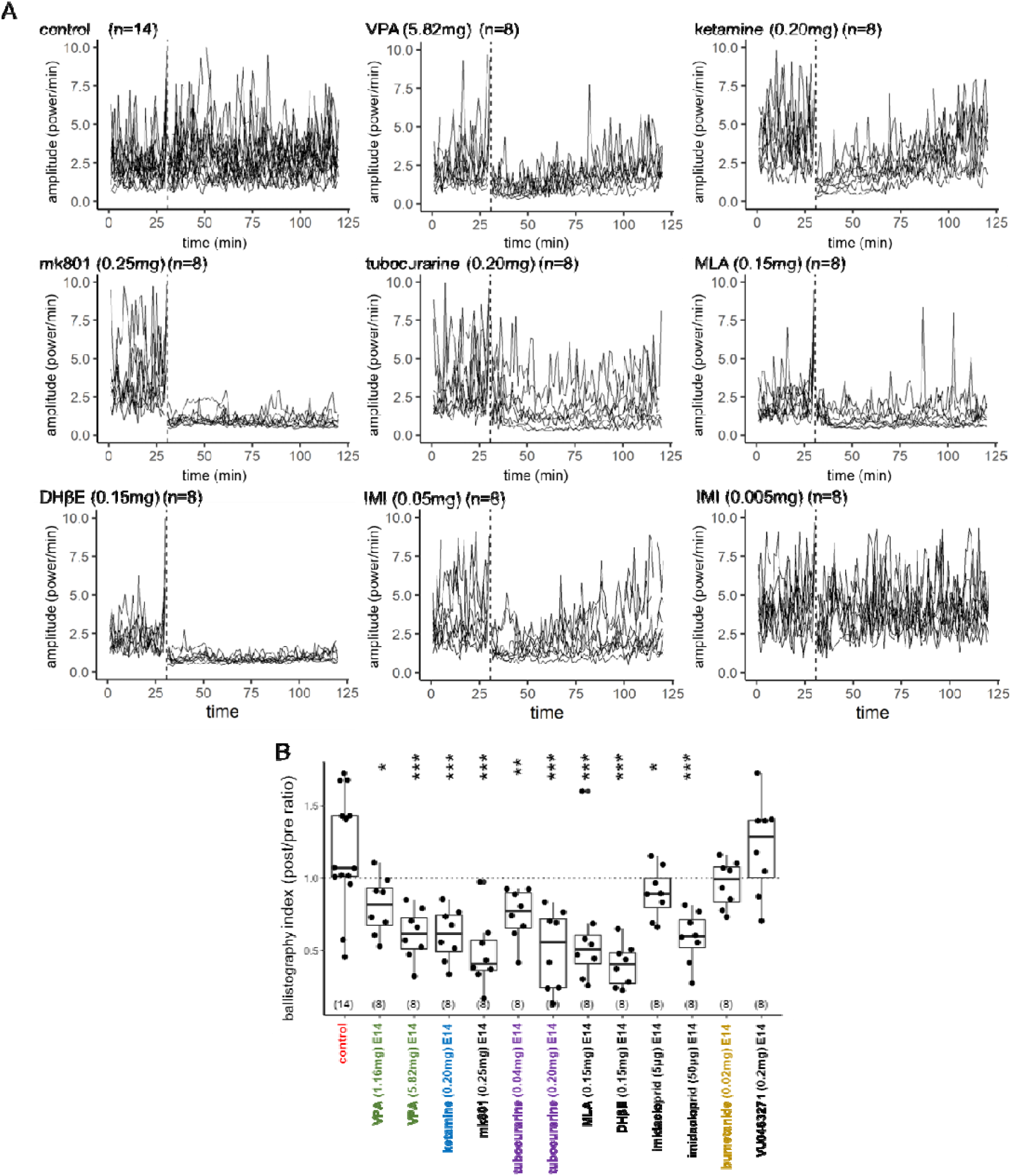

#### 1.3. Apparatus and procedures of imprinting and tests (experiment 2 and 3)

An I-shaped maze (10 cm wide, 70 cm long) was equipped with a 50-cm-long treadmill consisting of a rubber belt at the center and an LCD monitor at each end (**Figure 2Aa**) and placed in a dark room at 27~28°C. See our previous reports for details (Miura et al., 2018, 2020; Takemura et al. 2018; reference #31,54,55). Two training sessions (1 hour each) were given at 1 hour interval. An infrared sensor and a transparent Plexiglass partition were placed at a point 10 cm from one monitor, and the other monitor was occluded by an opaque partition. A loudspeaker behind the LCD monitor emitted mechanical sound synchronized with the toy movement on the screen. When chicks approached the monitor and hit the sensor, the rubber belt of the treadmill moved for a short period of 1.0 s, drawing the chick backward by about 30 cm at a time with the shortest intervals set at 0.1~2 sec. The apparatus was controlled by Arduino^®^. The treadmill motions were digitally counted, and the number of approaches was recorded for each chick. We monitored the chick behavior via a video camera set at the ceiling.

After 30 min to 1 hour pause period in a dark chamber kept at ~30°C, the trained chicks were tested in binary choice using the same apparatus without treadmill motions. In the BM test, one monitor displayed a linear point-light animation and the other monitor a walking one, both composed of white lights. In the subsequent imprinting test, one monitor displayed red toy (familiar object) and another yellow (novel object), both unaccompanied by the mechanical sound used in training. We consider that the subject chose one side when the whole body was located close (<30 cm) to the monitor (delineated by dashed line). Five-min tests were repeated twice after swapping the side, and the initial side was also counterbalanced among individuals. Tests were video-recorded. Intervals between test trials were set at 90-120sec. The difference in stay times gave the choice score in sec thus ranging in −600 to +600 sec.

In experiment 2, chicks were trained and tested on post-hatch 1 day (P1, at 12~36 hours after hatch) by the procedure shown in **Figure 2Ab**. In experiment 3, chicks were similarly trained and tested for the BM preference on P1, then re-trained by merged Lp(red) and Wp(yellow) on P2 (**Figure 3A**); the chicks were simultaneously exposed to two point-light animations, one walking in yellow Wp(yellow) and another linear in red Lp(red). Our recent study (Miura et al. 2020; reference #31) revealed that chicks exposed to the merged animations showed significantly higher preference to Wp[yellow] over Lp[red], if the BM preference of the chicks had been induced by the training using the red toy on P1.

#### 1.4. Video clips and point-light animations for training and tests

Two video clips of a rotating toy and 3 point-light animations were prepared as in our previous reports (Miura and Matsushima 2016, Miura et al. 2018, 2020; reference #30,31,54); see below for the list. Video clips were displayed on black background at a speed of 30 frames/s. We made editing by Adobe Premiere (Elements 7) and the color was set either to red (R: 255, G: 0, B: 0) or yellow (R: 255, G: 255, B: 0). These stimuli were displayed on the LCD monitors (size 10.4”, 800 × 600 pixels, Logitec LCM-T102AS, Japan; flash rate: 56–75 Hz, brightness: 230 cd/m2, pitch size: 0.264 × 0.264 mm) using free viewer software (A-player, version 6.0) on PC. The width of the presentation was set at 10.5 cm on the monitor, the surface of which was placed at 2.0 cm away from the terminal window of the maze.

|  |  |
| --- | --- |
| Moving-toy(red).wmv | P1 training and imprinting test (experiment 2,4) |
| Moving-toy(yellow).wmv | imprinting test (experiment 2,4) |
| Wp(white).wmv | BM test (experiment 2,4) |
| Lp(white).wmv | BM test (experiment 2,4) |
| Wp(yellow)-Lp(red).wmv | P2 training (experiment 4) |
| Wp(yellow).wmv | imprinting_1 (biological) test (experiment 4) |
| Lp(red).wmv | imprinting_1 (biological) test (experiment 4) |

#### 1.5. Chemical agents

Eggs (embryonic 14 days, E14, if not stated otherwise) received single injection of 200μL solution to the air chamber through one of two holes on the round edge of the shell; another hole served as air vent for smooth injection. The holes were sealed by mending tape. For each agent, the highest dosage was determined which (1) did not reduce the rate of hatching and (2) significantly suppressed the spontaneous fetal motions, then 1/5 diluted solution was tested. Control eggs received injection of the same amount of vehicle, distilled water.

Name, code and company of the chemical agents

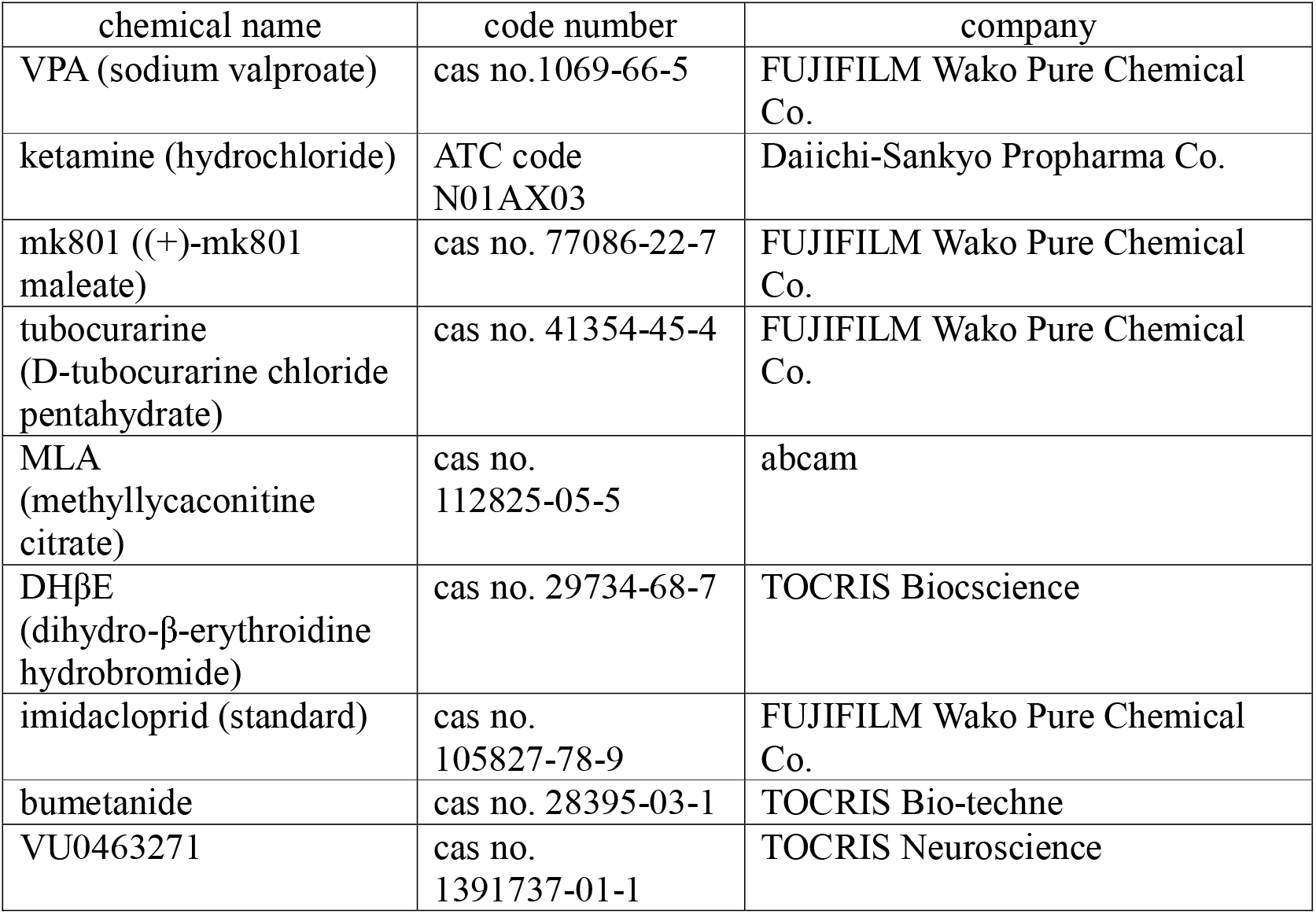

Amount of agents injected / egg ~ 50g

| chemical agent | molecular weight | µmole |
| --- | --- | --- |
| VPA 5.82mg | 166.19 | 35.0 |
| ketamine 0.20mg | 274.19 | 0.729 |
| mk801 0.25mg | 337.37 | 0.741 |
| tubocurarine 0.20mg | 771.72 | 0.259 |
| MLA 0.15mg | 874.92 | 0.171 |
| DHβE 0.15mg | 356.26 | 0.421 |
| imidacloprid 0.05mg | 255.66 | 0.196 |
| bumetanide 0.02mg | 364.42 | 0.0549 |
| VU0463271 0.2mg | 387 | 0.517 |

#### 1.6. Histone acetylation assay in cultured brain cells (experiment 4)

Embryos (E7 and E14) were aseptically harvested from eggs and brain tissues were separated. Brain tissues were finely minced with scissors in sterile phosphate-buffered saline (PBS) and dissociated by pipetting with micropipette tips. Dissociated brain tissues were suspended in a brain culture medium composed of DMEM/F12 (048-29785, FUJIFILM Wako Pure Chemical Co., Tokyo, Japan), 10% foetal bovine serum (S-FBS-NL-015, Serana Europe GmbH, Brandenburg, Germany), 1x Antibiotic-Antimycotic (15240-096, Life Technologies Co., CA, USA), and maintained on coverslip glasses coated with collagen type I-C (631-00771, FUJIFILM Wako Pure Chemical Co., Tokyo, Japan) for 4 days at 37°C in 5% CO_2_ condition.

The cultured brain tissues were incubated with ketamine (66 or 6.6 μg/mL), VPA (1.2 mM or 0.12 mM) or vehicle PBS in a brain culture medium without FBS for 2 hours at 37°C in 5% CO_2_. After incubation, cultured brain tissues were fixed with 4% paraformaldehyde in PBS for 5 mins, blocked with a blocking solution (4% normal goat serum (S-1000, VECTOR laboratories Inc., CA, USA), 1% blocking reagent (1096176, Roche Ltd., Basel, Swiss) and 0.5% Tween20 in PBS) for 30 mins, and then incubated with 1:3000 diluted Anti-acetyl Histone H3 (Lys27), mouse monoclonal antibody (MA309A, TaKaRa Bio Inc., Shiga, Japan) in the blocking solution for 16 hours at 4□in a humidity chamber. Following the first antibody incubation, tissues were incubated with 1:2000 diluted Goat anti-Mouse IgG, Alexa Fluor 555 antibody (A21422, ThermoFisher Scientific, MA, USA) in the blocking solution for 2 hours at room temperature in a humidity chamber. Stained tissues were mounted in Mounting Medium with DAPI (H-1200, VECTOR Laboratories Inc.) and imaged using a BZ-X810 microscope (Keyence Co., Osaka, Japan). In each duplicated experimental group, three fields of view were taken per coverslip, and the acetylation level of H3K27 in the nuclei of 100 randomly selected cells from six fields of view was measured as fluorescence intensity using Image J.

#### 1.7. Brain weight and isotropic fractionation measurement of brain cells (experiment 5 and 6)

The P1 chicks were transcardially perfused by 4% paraformaldehyde in 0.1M phosphate buffer under a deep anesthesia by i.m. injection of 0.8mL ketamine-xylazine cocktail; a 1:1 mixture of 10mg/mL ketamine hydrochloride (Daiichi-Sankyo Propharma Co.) and 2mg/mL xylazine (Sigma-Aldrich Co.).

The whole brain (cut at the caudal end of medulla oblongata) was dissected and post-fixed in the fixative for overnight, stored for 2~4 days in phosphate-buffered saline (PBS) at 4°C and weighed. The telencephalon was isolate at the rostral level of diencephalon and manually chopped in small pieces by scalpel. The tissue was weighed and homogenized in a detergent solution (40mM sodium citrate and 1% Triton X-100) using a 7mL glass Tenbroeck tissue homogenizer. The homogenate and several washes of the homogenizer were transferred to a 50ml falcon tube using a glass pipette. To visualize cell nuclei DAPI (4,6-diamidino-2-phenylindoledihydrochloride, Invitrogen, Carlsbad, Calif., USA) was added to the suspension from a stock solution at 20 mg/L (dilution 1:20 to 1:50) and the final volume of the suspension was recorded. To estimate the total amount of cells in the suspension, the nuclear density was quantified in a Neubauer chamber by a fluorescent microscope (BX-50, Olympus Co., Tokyo, Japan) according to the protocol developed by Herculano-Houzel and Lent (2005).

To estimate the number of matured neuron, aliquot (500 μL) of the nuclear suspension washed with PB (0.1M) and incubated overnight in the dark at 4°C with Cy3-labeled rabbit polyclonal neuronal nuclear antigen antibody (1:150 dilution; NeuN, RRID:AB_11204707). The percentage of neuronal nuceli was determined by counting at least 500 DAPI-stained nuclei and establishing the fraction that was also NeuN positive, according to the protocol developed by Herculano-Houzel and Lent (2005). Briefly, the total number of neurons in each sample was determined by multiplying the total number of cells in the structure by the NeuN-positive fraction obtained. The total number of non-neuronal cells was determined by subtracting the total number of neurons from the total number of cells. Densities of neurons and non-neurons (cell/mg) were determined by dividing the number of neurons or non-neurons by the mass (mg) of the structure.

### 2. Statistical analyses

Statistical computations were performed on RStudio version 3.6.3 (2020-02-29). For single and multiple regression analyses, linear models were constructed using the function *lm()* with the control group as the reference set; one model was calculated for each set of experimental groups. Multiple regression including variables such as *sex*, *batch* and *run* (locomotor counts during training) appeared not to improve the statistical judgements; we therefore adopted single regression by *drug* (and dose) in most cases.

In experiment 6, where gene expression profiles were examined, IMM and E14 served the reference for the brain region and the age, respectively. Significance is coded as 0 < *** < 0.001 < ** < 0.01 < * <0.05, and ns means p-value ≥ 0.05.

#### 2.1. Experiment 1

##### 2.1.1. Ballistography index explained by drug (Figure 1C)

fit_exp_1_ballistography_selected <- lm(ratio ~ label, data = dataset_exp_1_ballistography_fig) summary(fit_exp_1_ballistography_selected)

Coefficients:

| label (drug) | estimate | std. error | t-value | Pr(>\|t\|) | signif. code |
| --- | --- | --- | --- | --- | --- |
| (intercept) | +1.18063 | 0.07899 | 14.947 | < 2e-16 | *** |
| VPA 5.82mg | -0.56805 | 0.13099 | -4.337 | 6.23e-05 | *** |
| ketamine 0.20mg | -0.57210 | 0.13099 | -4.368 | 5.60e-05 | *** |
| mk801 0.25mg | -0.69986 | 0.13099 | -5.343 | 1.80e-06 | *** |
| tubocurarine 0.20mg | -0.67630 | 0.13099 | -5.163 | 3.46e-06 | *** |
| MLA 0.15mg | -0.56939 | 0.13099 | -4.347 | 6.01e-05 | *** |
| DHβE 0.15mg | -0.78066 | 0.13099 | -5.960 | 1.87e-07 | *** |

residual standard error = 0.2956 (Df = 55)
multiple r-squared = 0.5006, adjusted r-squared = 0.4461
F-statistic = 9.188 (Df = 6, 55), p-value = 5.765e-07

#### 2.2 Experiment 2, VPA, ketamine and tubocurarine (Figure 2B)

##### 2.2.1. BM score explained by drug (Figure 2Ba)

fit_exp2_bm_selected <- lm (bm ~ label, data = dataset_exp_2_bm_imprint_fig) summary (fit_exp2_bm_selected)

Coefficients:

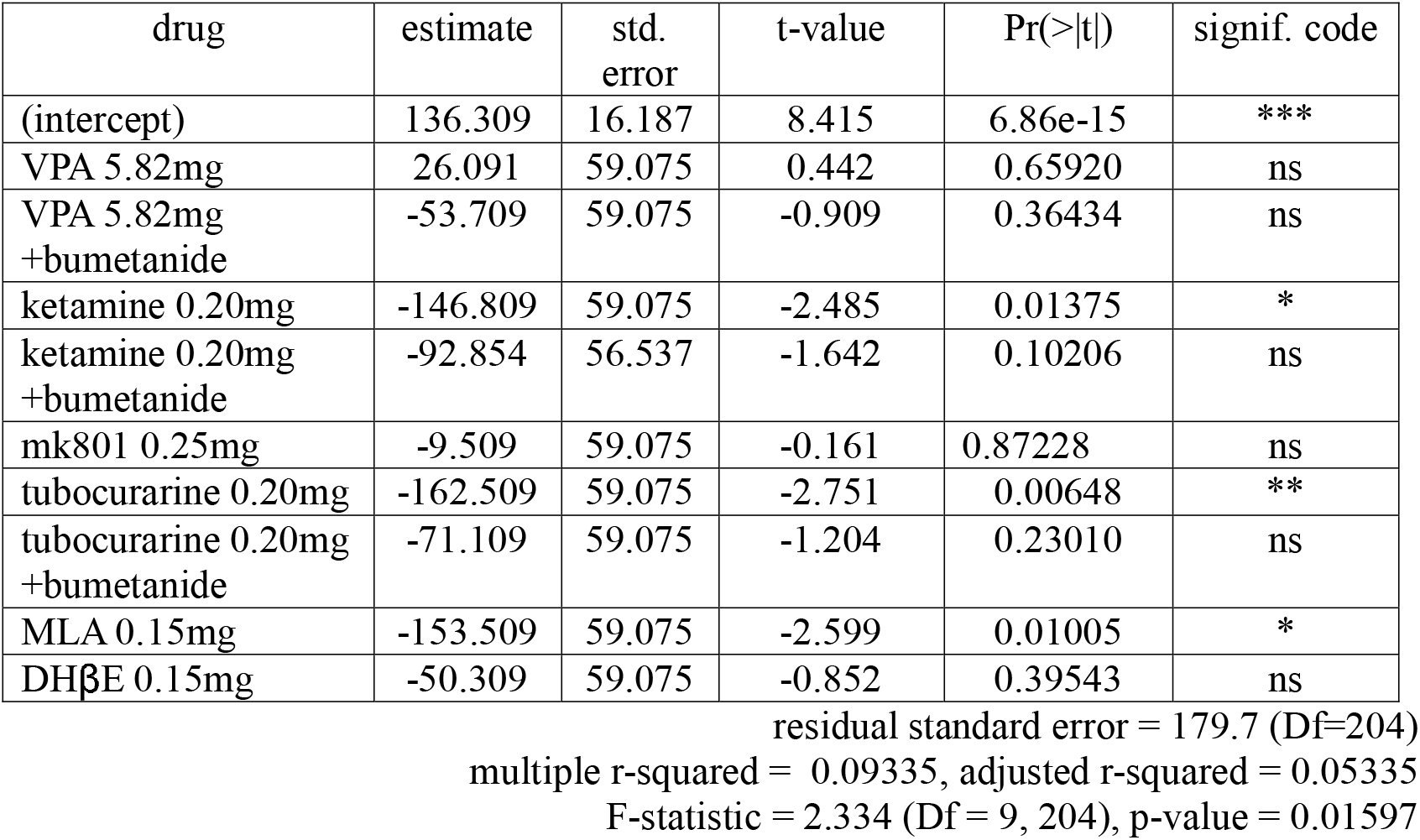

| drug | estimate | std. error | t-value | Pr(>\|t\|) | signif. code |
| --- | --- | --- | --- | --- | --- |
| (intercept) | 136.309 | 16.187 | 8.415 | 6.86e-15 | *** |
| VPA 5.82mg | 26.091 | 59.075 | 0.442 | 0.65920 | ns |
| VPA 5.82mg<br>+bumetanide | -53.709 | 59.075 | -0.909 | 0.36434 | ns |
| ketamine 0.20mg | -146.809 | 59.075 | -2.485 | 0.01375 | * |
| ketamine 0.20mg<br>+bumetanide | -92.854 | 56.537 | -1.642 | 0.10206 | ns |
| mk801 0.25mg | -9.509 | 59.075 | -0.161 | 0.87228 | ns |
| tubocurarine 0.20mg | -162.509 | 59.075 | -2.751 | 0.00648 | ** |
| tubocurarine 0.20mg<br>+bumetanide | -71.109 | 59.075 | -1.204 | 0.23010 | ns |
| MLA 0.15mg | -153.509 | 59.075 | -2.599 | 0.01005 | * |
| DHβE 0.15mg | -50.309 | 59.075 | -0.852 | 0.39543 | ns |

residual standard error = 179.7 (Df=204)
multiple r-squared = 0.09335, adjusted r-squared = 0.05335
F-statistic = 2.334 (Df = 9, 204), p-value = 0.01597

##### 2.2.2. imprinting score explained by drug (Figure 2Bb)

fit_exp2_imprint_selected <- lm (imprint ~ label, data = dataset_exp_2_bm_imprint_fig) summary (fit_exp2_imprint_selected)

Coefficients:

| drug | estimate | std. error | t-value | Pr(>\|t\|) | signif. code |
| --- | --- | --- | --- | --- | --- |
| (intercept) | 419.52 | 19.80 | 21.183 | < 2e-16 | *** |
| VPA 5.82mg | -316.22 | 72.22 | -4.378 | 1.91e-05 | *** |
| VPA 5.82mg<br>+bumetanide | 84.68 | 72.22 | 1.172 | 0.2424 | ns |
| ketamine 0.20mg | -118.62 | 72.22 | -1.642 | 0.1020 | ns |
| ketamine 0.20mg<br>+bumetanide | -26.07 | 69.12 | -0.377 | 0.7065 | ns |
| mk801 0.25mg | 3.88 | 72.22 | 0.054 | 0.9572 | ns |
| tubocurarine 0.20mg | 47.08 | 72.22 | 0.652 | 0.5152 | ns |
| tubocurarine 0.20mg<br>+bumetanide | -5.42 | 72.22 | -0.075 | 0.9402 | ns |
| MLA 0.15mg | -160.32 | 72.22 | -2.220 | 0.0275 | * |
| DHβE 0.15mg | -151.02 | 72.22 | -2.091 | 0.0378 | * |

residual standard error = 219.6 (DF = 204) multiple r-squared = 0.1348, adjusted r-squared = 0.09664 F-statistic = 3.532 (DF = 9, 204), p-value = 0.0004322

#### 2.3. Experiment 2, imidacloprid (Figure 2C)

##### 2.3.1. BM score explained by drug (Figure 2Ca)

fit_exp_3_bm <- lm (bm ~ label, data = dataset_exp_2_bm_imprint_ctrl_imi) summary (fit_exp_3_bm)

Coefficients:

| drug | estimate | std. error | t-value | Pr(>\|t\|) | signif. code |
| --- | --- | --- | --- | --- | --- |
| (intercept) | 117.66 | 21.05 | 5.589 | 1.42e-07 | *** |
| E0 IMI 0.005mg | -28.25 | 47.90 | -0.590 | 0.5564 | ns |
| E0 IMI 0.05mg | -109.98 | 45.82 | -2.400 | 0.0179 | * |
| E14 IMI 0.005mg | -114.16 | 59.92 | -1.905 | 0.0591 | ns |
| E14 IMI 0.05mg | -137.06 | 59.92 | -2.288 | 0.0239 | * |

residual standard error = 177.4 (DF = 122) multiple r-squared = 0.08534, adjusted r-squared = 0.05535 F-statistic = 2.846 (DF = 4, 122), p-value = 0.02689

##### 2.3.2. imprinting score explained by drug (Figure 2Cb)

fit_exp_3_imprint <- lm (imprint ~ label, data = dataset_exp_2_bm_imprint_ctrl_imi) summary (fit_exp_3_imprint)

Coefficients:

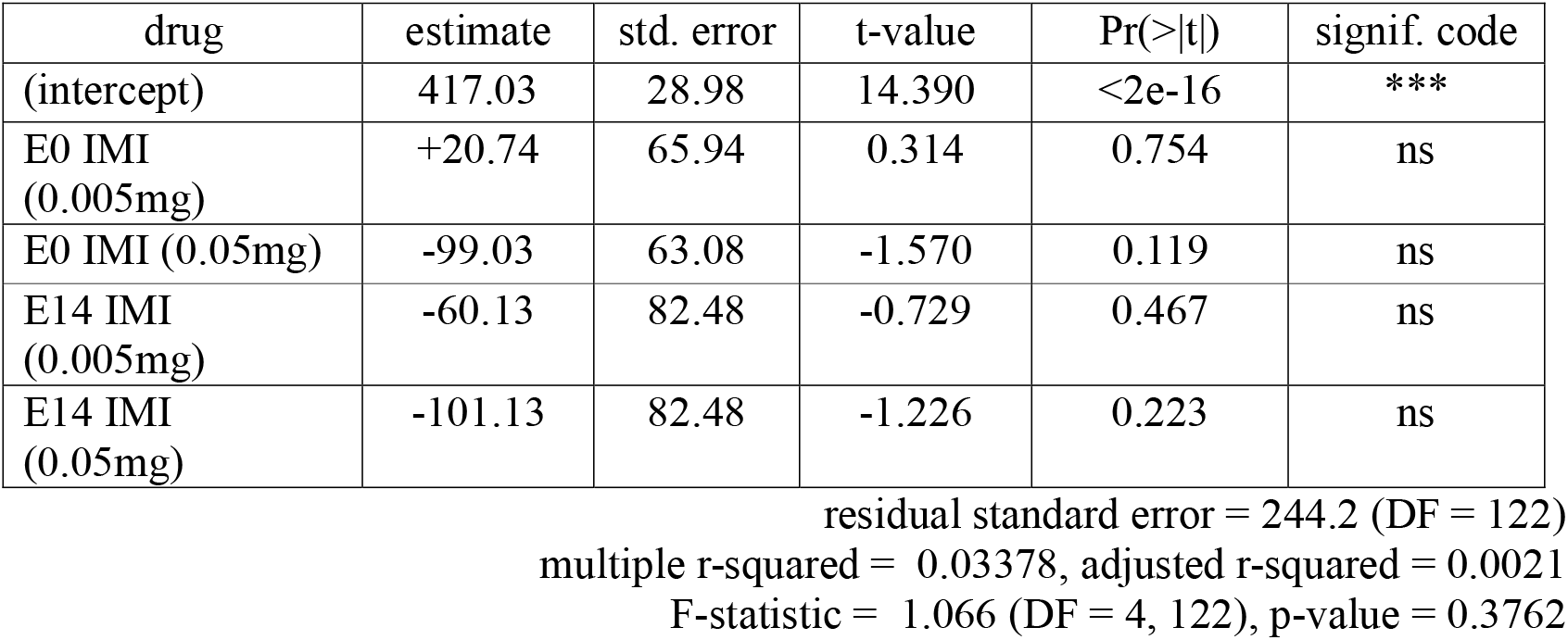

| drug | estimate | std. error | t-value | Pr(>\|t\|) | signif. code |
| --- | --- | --- | --- | --- | --- |
| (intercept) | 417.03 | 28.98 | 14.390 | <2e-16 | *** |
| E0 IMI<br>(0.005mg) | +20.74 | 65.94 | 0.314 | 0.754 | ns |
| E0 IMI (0.05mg) | -99.03 | 63.08 | -1.570 | 0.119 | ns |
| E14 IMI<br>(0.005mg) | -60.13 | 82.48 | -0.729 | 0.467 | ns |
| E14 IMI<br>(0.05mg) | -101.13 | 82.48 | -1.226 | 0.223 | ns |

residual standard error = 244.2 (DF = 122) multiple r-squared = 0.03378, adjusted r-squared = 0.0021 F-statistic = 1.066 (DF = 4, 122), p-value = 0.3762

#### 2.4. Bootstrapping analysis of Experiment 2

We made post-hoc bootstrapping analysis of the control data (123 chicks) to obtain the distribution (probability density) of the mean of samples composed of n=10 chicks. For each of BM and imprinting scores, 10 chicks were randomly sampled 10,000 times from the control data. For each sample from 10 chicks, difference between the sample average (n=10) and the average of the unsampled chicks (n=113) were calculated, thus yielding a set of 10,000 values. Distribution of the difference value was represented as probability density curves shown below (**Figure S2**), and the critical points where the area below the curve occupied 95% and 99% were obtained by using *quantile()*.

**Figure S2.**
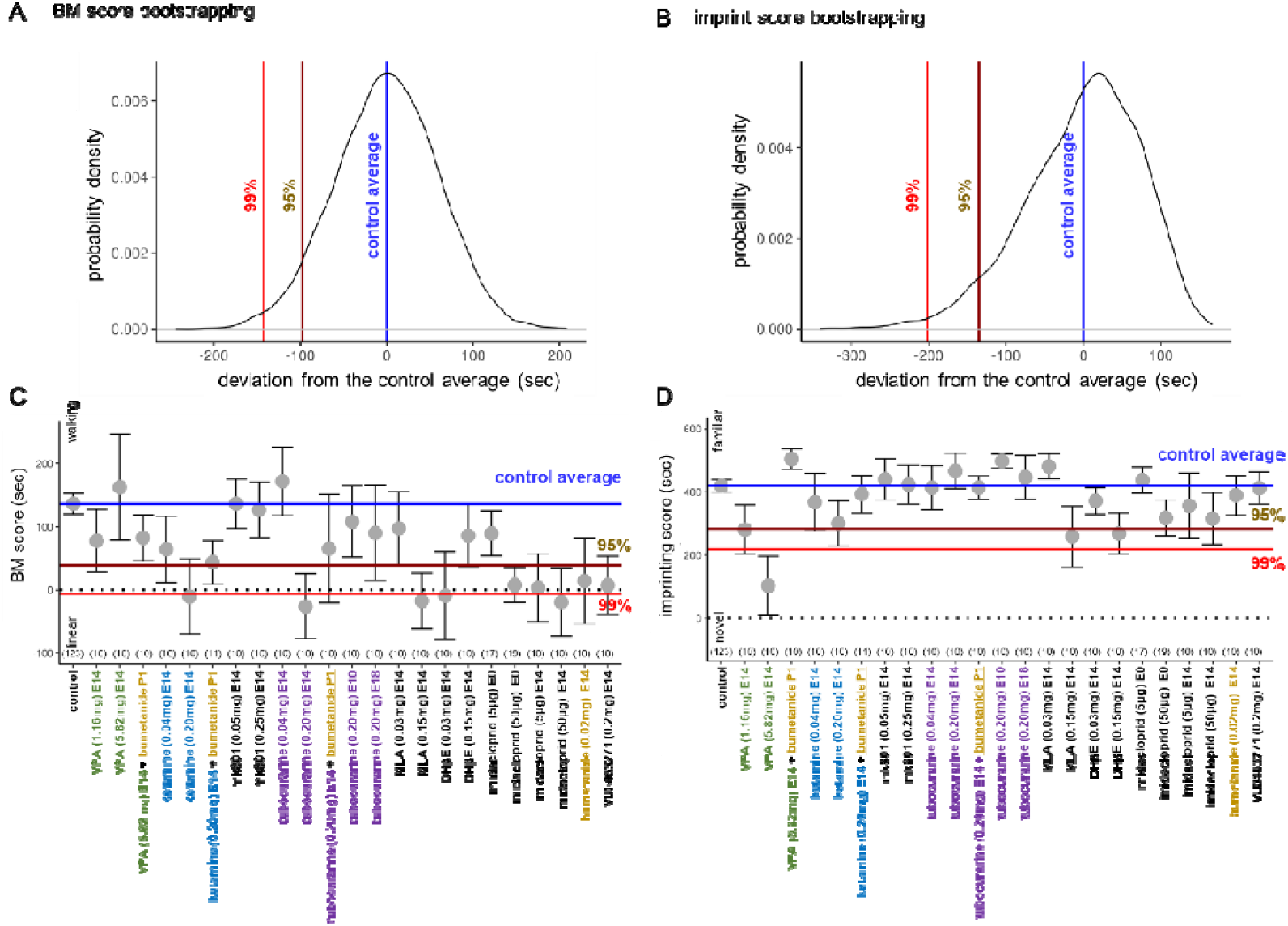

#### 2.5. Experiment 3 (Figure 3)

##### 2.5.1. BM score explained by drug (Figure 3B)

fit_exp4_bm_drug <- lm(bm ~ drug, data = dataset_exp_4_double_imprint) summary (fit_exp4_bm_drug)

Coefficients:

| drug | estimate | std. error | t-value | Pr(>\|t\|) | signif. code |
| --- | --- | --- | --- | --- | --- |
| control | 145.80 | 52.24 | 2.791 | 0.00781 | ** |
| VPA 5.82mg | -18.22 | 78.37 | -0.232 | 0.81729 | ns |
| ketamine 0.20mg | -172.30 | 82.61 | -2.086 | 0.04296 | * |
| tubocurarine 0.20mg | -235.00 | 82.61 | -2.845 | 0.00678 | ** |

residual standard error = 202.3 (DF = 43)
multiple r-squared = 0.2076, adjusted r-squared = 0.1523
F-statistic = 3.755 (DF = 3, 43), p-value = 0.01758

##### 2.5.2. imprinting_1 (biological) score explained by drug (Figure 3Ca)

fit_exp4_imprint_1 <- lm(imprint_1 ~ drug, data = dataset_exp_4_double_imprint) summary (fit_exp4_imprint_1)

Coefficients:

| drug | estimate | std. error | t-value | Pr(>\|t\|) | signif. code |
| --- | --- | --- | --- | --- | --- |
| control | 37.13 | 59.77 | 0.621 | 0.5377 | ns |
| VPA 5.82mg | -213.05 | 89.66 | -2.376 | 0.0220 | * |
| ketamine 0.20mg | -201.53 | 94.51 | -2.132 | 0.0387 | * |
| tubocurarine 0.20mg | -60.33 | 94.51 | -0.638 | 0.5266 | ns |

residual standard error = 231.5 (DF = 43) multiple r-squared = 0.1533, adjusted r-squared = 0.09428 F-statistic = 2.596 (DF = 3, 43), p-value = 0.06465

##### 2.5.3. imprinting_1 (biological) score explained by drug and BM score (Figure 3Cb)

fit_exp4_imprint_1_bm <- lm(imprint_1 ~ drug * bm, data = dataset_exp_4_double_imprint) summary (fit_exp4_imprint_1_bm)

Coefficients:

| drug/BM | estimate | std. error | t-value | Pr(>\|t\|) | signif. code |
| --- | --- | --- | --- | --- | --- |
| (intercept) | -128.6864 | 72.0850 | -1.785 | 0.082007 | ns |
| VPA 5.82mg | -1.7390 | 99.4170 | -0.017 | 0.986134 | ns |
| ketamine 0.20mg | -36.8073 | 98.1233 | -0.375 | 0.709609 | ns |
| tubocurarine 0.20mg | 104.6133 | 103.3479 | 1.012 | 0.317660 | ns |
| BM | 1.1373 | 0.3286 | 3.462 | 0.001317 | ** |
| VPA*BM | -1.4939 | 0.4161 | -3.591 | 0.000911 | *** |
| ketamine*BM | -1.1786 | 0.4706 | -2.505 | 0.016545 | * |
| tubocurarine*BM | -1.1471 | 0.5003 | -2.293 | 0.027323 | * |

residual standard error = 208.6 (DF = 39) multiple r-squared = 0.3764, adjusted r-squared = 0.2645 F-statistic = 3.363 (DF = 7, 39), p-value = 0.00668

##### 2.5.4. imprinting_2 (artifact) score explained by drug (Figure 3Da)

fit_exp4_imprint_2 <- lm(imprint_2 ~ drug, data = dataset_exp_4_double_imprint) summary (fit_exp4_imprint_2)

Coefficients:

| drug | estimate | std. error | t-value | Pr(>\|t\|) | signif. code |
| --- | --- | --- | --- | --- | --- |
| control | 487.13 | 68.66 | 7.095 | 9.38e-09 | *** |
| VPA 5.82mg | -277.38 | 102.99 | -2.693 | 0.0100 | * |
| ketamine 0.20mg | -194.93 | 108.56 | -1.796 | 0.0796 | ns |
| tubocurarine 0.20mg | -91.23 | 108.56 | -0.840 | 0.4053 | ns |

residual standard error = 265.9 (DF = 43) multiple r-squared = 0.1582, adjusted r-squared = 0.09942 F-statistic = 2.693 (DF = 3, 43), p-value = 0.0579

##### 2.5.5. imprinting_2 (artifact) score explained by drug and BM (Figure 3Db)

fit_exp4_imprint_2_bm <- lm(imprint_2 ~ drug*bm, data = dataset_exp_4_double_imprint) summary (fit_exp4_imprint_2_bm)

Coefficients:

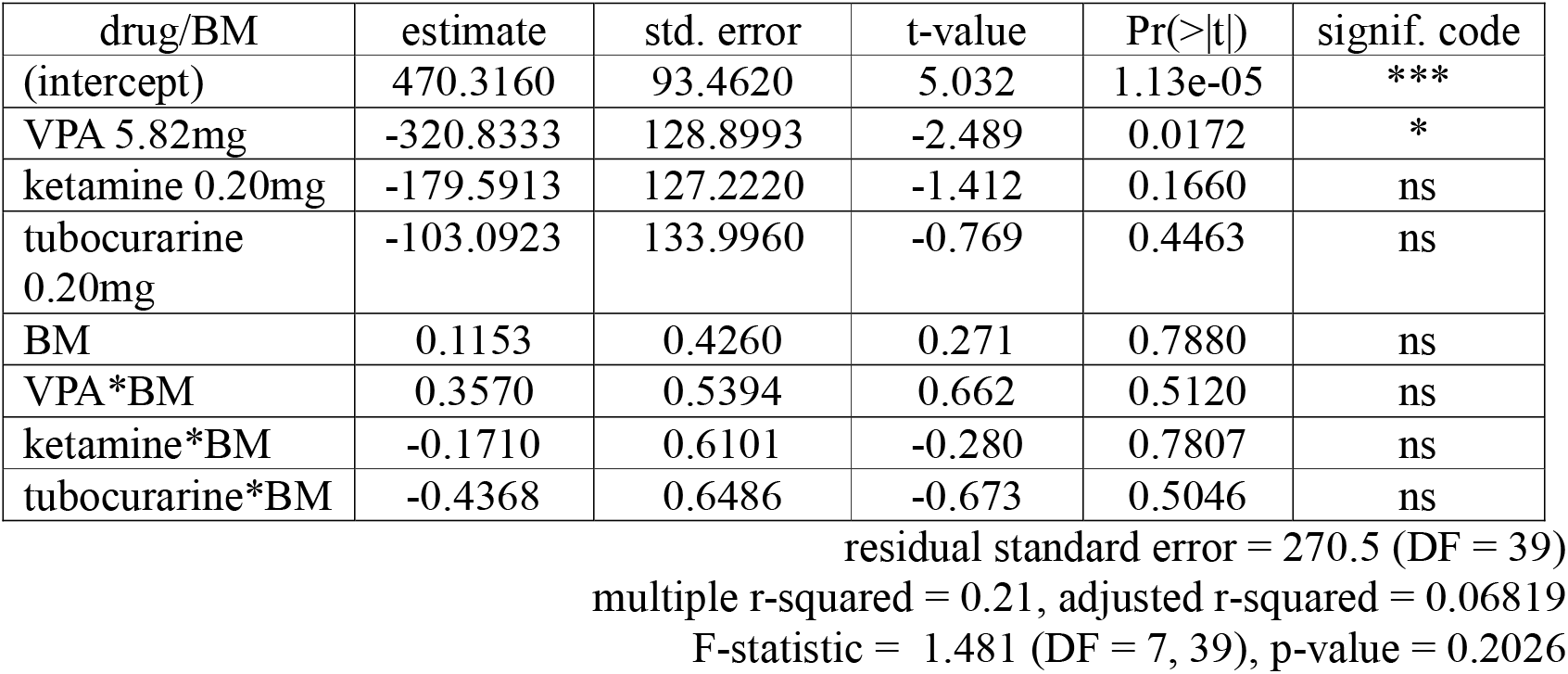

#### 2.6. Experiment 4 (Figure 4A) H3K27ac fluorescence explained by drug

fit_exp_5_h3k27ac <- lm(flu ~ drug, data = dataset_exp_5_h3k27ac) summary(fit_exp_5_h3k27ac)

Coefficients:

| drug | estimate | std. error | t-value | Pr(>\|t\|) | signif. code |
| --- | --- | --- | --- | --- | --- |
| (intercept) | +7.4293 | 0.3156 | 23.537 | <2e-16 | *** |
| VPA 0.12mM | +0.5917 | 0.4464 | 1.326 | 0.186 | ns |
| VPA 1.2mM | +11.9921 | 0.4464 | 26.865 | <2e-16 | *** |
| ketamine 6.6<br>µg/mL | □<br>0.4920 | 0.4464 | □ 1.102 | 0.271 | ns |
| ketamine 66<br>µg/mL | +0.0230 | 0.4464 | 0.052 | 0.959 | ns |

residual standard error = 3.156 (DF = 495)
multiple r-squared = 0.7, adjusted r-squared = 0.6975
F-statistic = 288.7 (DF=4, 495), p-value: < 2.2e-16

#### 2.7. Experiment 5 and 6 (Figure 4B,C)

##### 2.7.1. brain weight (Figure 4Ba)

fit_exp_6_abs_w_brain <- lm (w_brain ~sex * drug, data = dataset_exp_6_brain_weight) summary (fit_exp_6_abs_w_brain)

Coefficients:

| drug/sex | estimate | std. error | t-value | Pr(>\|t\|) | signif. code |
| --- | --- | --- | --- | --- | --- |
| (intercept) | 1.00174 | 0.01520 | 65.887 | < 2e-16 | *** |
| sex | 0.03101 | 0.02135 | 1.453 | 0.148404 | ns |
| VPA 5.82mg | -0.08410 | 0.02426 | -3.467 | 0.000683 | *** |
| ketamine 0.20mg | -0.04889 | 0.02891 | -1.691 | 0.092845 | ns |
| tubocurarine | -0.03094 | 0.04246 | -0.729 | 0.467397 | ns |
| 0.20mg |  |  |  |  |  |
| sex* VPA | 0.02942 | 0.03266 | 0.901 | 0.369059 | ns |
| sex* ketamine | 0.01340 | 0.03939 | 0.340 | 0.734245 | ns |
| sex* tubocurarine | 0.02419 | 0.06000 | 0.403 | 0.687336 | ns |

residual standard error = 0.08865 (DF = 153) multiple r-squared = 0.1514, adjusted r-squared = 0.1126 F-statistic= 3.901 (Df = 7, 15), p-value = 0.0006117

##### 2.7.2. body weight (Figure 4Bb)

fit_exp_6_abs_w_body <- lm (w_body ~ sex * drug, data = dataset_exp_6_brain_weight) summary (fit_exp_6_abs_w_body)

Coefficients:

| drug/sex | estimate | std. error | t-value | Pr(>\|t\|) | signif. code |
| --- | --- | --- | --- | --- | --- |
| (intercept) | 39.65882 | 0.54955 | 72.166 | <2e-16 | *** |
| sex | 0.17832 | 0.87678 | 0.231 | 0.818 | ns |
| VPA 5.82mg | -1.3451 | 0.02426 | -1.534 | 0.127 | ns |
| ketamine 0.20mg | -0.06652 | 0.02891 | -0.064 | 0.949 | ns |
| tubocurarine<br>0.20mg | 1.10118 | 1.53481 | 0.717 | 0.474 | ns |
| sex* VPA | 0.85643 | 1.18040 | 0.726 | 0.469 | ns |
| sex* ketamine | -1.36438 | 1.42373 | -0.958 | 0.339 | ns |
| sex* tubocurarine | -1.91832 | 2.16856 | -0.885 | 0.378 | ns |

Residual standard error = 3.204 (DF = 153) multiple r-squared = 0.03717, adjusted r-squared = -0.006877 F-statistic= 0.8439 (Df = 7, 153), p-value = 0.5527

##### 2.7.3. NeuN-positive cell ratio (Figure 4Ca)

fit_exp_6_neuron_glia_ratio <- lm(neuron_ratio ~ sex * drug, data = dataset_exp_6_neuron_glia) summary (fit_exp_6_neuron_glia_ratio)

Coefficients:

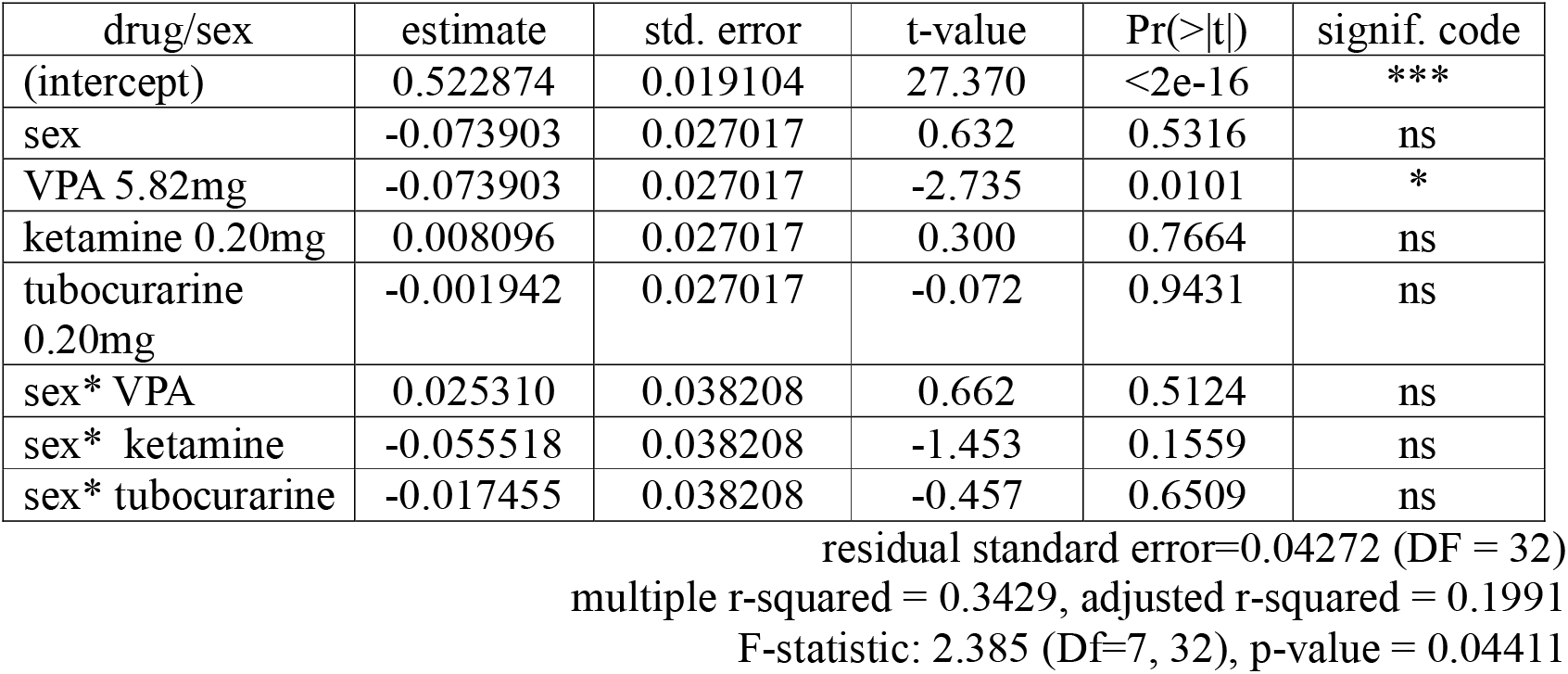

##### 2.7.4. total cell number (Figure 4Cb)

fit_exp_6_cell_count <- lm(cell_number ~ sex * drug, data = dataset_exp_6_neuron_glia) summary (fit_exp_6_cell_count)

Coefficients:

| drug/sex | estimate | std. error | t-value | Pr(>\|t\|) | signif. code |
| --- | --- | --- | --- | --- | --- |
| (intercept) | 32.806 | 3.128 | 10.488 | 6.97e-12 | *** |
| sex | 3.626 | 4.424 | 0.820 | 0.418 | ns |
| VPA 5.82mg | -0.599 | 4.424 | -0.135 | 0.893 | ns |
| ketamine 0.20mg | -1.412 | 4.424 | -0.319 | 0.752 | ns |
| tubocurarine<br>0.20mg | 2.088 | 4.424 | 0.472 | 0.640 | ns |
| sex* VPA | -2.719 | 6.256 | -0.435 | 0.667 | ns |
| sex* ketamine | -2.004 | 6.256 | -0.320 | 0.751 | ns |
| sex* tubocurarine | -5.436 | 6.256 | -0.869 | 0.391 | ns |

residual standard error=6.994 (DF = 32)
multiple r-squared = 0.05307, adjusted r-squared = -0.1541
F-statistic= 0.2562 (Df = 7, 32), p-value = 0.9663

